# Concurrent spinal and brain imaging with optically pumped magnetometers

**DOI:** 10.1101/2022.05.12.491623

**Authors:** Lydia C. Mardell, George C. O’Neill, Tim M. Tierney, Ryan C. Timms, Catharina Zich, Gareth R. Barnes, Sven Bestmann

## Abstract

The spinal cord and its interactions with the brain are fundamental for movement control and somatosensation. However, brain and spinal cord electrophysiology in humans have largely been treated as distinct enterprises, in part due to the relative inaccessibility of the spinal cord. Consequently, there is a dearth of knowledge on human spinal electrophysiology, including the multiple pathologies of the central nervous system that affect the spinal cord as well as the brain. Here we exploit recent advances in the development of wearable optically pumped magnetometers (OPMs) which can be flexibly arranged to provide coverage of both the spinal cord and the brain concurrently in unconstrained environments. Our system for magnetospinoencephalography (MSEG) measures both spinal and cortical signals simultaneously by employing a custom-made spinal scanning cast. We evidence the utility of such a system by recording simultaneous spinal and cortical evoked responses to median nerve stimulation, demonstrating the novel ability for concurrent non-invasive millisecond imaging of brain and spinal cord.

## Introduction

The spinal cord transmits sensory information from the periphery to the brain and constitutes the final stage of processing within the central nervous system (CNS) for the production of movement. Yet in humans, it is also one of the most difficult structures of the central nervous system to study^1–4^.

One fundamental barrier to our understanding of human spinal cord function in health and disease is the limited ability to study its electrophysiology, due to the inaccessibility of the spinal cord^4^. Consequently, there is an unmet need for precise spinal imaging with millisecond precision because many acquired and neurodegenerative diseases that act on the brain also affect the spinal cord, and vice versa^5–8^. For example, acute and chronic pain provoke both spinal and cortical changes^9–11^, damage to the spinal cord from spinal cord injury leads to profound retrograde degeneration^8^, and damage to the brain after stroke can result in anterograde changes at spinal levels^12,13^. Yet in humans little is known about the neuropathology of the spinal cord and cortico-spinal interactions.

One implication is that our understanding of many neurological disorders is largely agnostic to their spinal mechanisms, and hence development of novel treatment approaches or rehabilitation regimes often overlook the role of the spinal cord or rely on small animal disease models^1^. Development of electrophysiological imaging of the human spinal cord, concurrently with imaging of the brain, will thus be foundational to deeper understanding of the spinal cord in health and disease.

### Current measures of spinal activity

Currently, precise recordings of human spinal electrophysiology are dominated by direct, invasive measurements, often recorded during surgery^14–16^. Commonly used non-invasive techniques for measuring spinal electrophysiology generally rely on surface electrodes, which, for example, can detect spinal cord evoked potentials (SCEPs) following electrical stimulation of peripheral nerves^17–20^. SCEPs recorded with skin and scalp electrodes, while non-invasive and easy to record, can be difficult to interpret due to the distortion of the volume currents caused to surrounding tissue which in turn limits precise estimates of the underlying sources^21^.

These limitations are overcome with magnetospinography (MSG) which uses superconducting quantum interference devices (SQUIDs) to detect the magnetic field generated by the spinal cord^22–26^. The magnetic fields detected with MSG are not as distorted by the tissue surrounding the spinal cord or muscle artefacts, meaning source analysis is more reliable^23,27^. Recent MSG systems are based upon custom-built arrays of SQUID sensors within rigid cryogenic vessels (dewars) filled with liquid helium^21^. Both reclining and supine position scanning systems have been developed, optimized to record from the cervical spine^22,28^. These systems allow the detection of magnetic fields generated by spinal neurons and innervating nerves, including early spinal cord evoked fields (SCEFs) following supramaximal peripheral nerve stimulation^21,28^.

SQUID based MSG, despite its significant achievements, does however have some limitations. It requires the participant to remain still^22^, making studies of spinal cord electrophysiology during movement, or studies of patient cohorts that have difficulty staying immobile, challenging. The coverage of the spinal cord is dictated by the size and shape of the generic fixed SQUID array rather fit to the subject’s anatomy. Finally, these systems currently cannot record simultaneously from the brain and spinal cord, preventing the study of brain-spinal interactions.

### Magnetospinoencephalography (MSEG) with optically pumped magnetometers

Here we present a novel system for concurrent spinal and brain recording with millisecond precision. To this end, we leverage the development of wearable magnetoencephalography (MEG) systems^29^ incorporating optically pumped magnetometers (OPMs). OPMs are sensing devices that can measure the absolute magnetic field due to neuronal current flow, which were previously commonly recorded with large, rigid superconducting sensors^30^. Recent progress in miniaturization and commercialization now provides access to OPMs that are lightweight and small, approximately the size of a LEGO brick (www.quspin.com). Usually, these sensors are placed in a helmet that can be worn by the participant^31–34^. In contrast to traditional SQUID-based MEG systems, which comprise a rigid superconducting flask, the field-sensitive volume of the device can be placed flexibly within 6 – 8 mm of the skin surface. This boosts signal magnitude (by the square of distance) and enables fundamentally better spatial resolution^35,36^. Flexible sensor placement and wearability of OPM-based MEG (OP-MEG) has enabled detection of neuromagnetic fields generated by deep sources such as the cerebellum^37^ and hippocampus^38,39^. OPMs thus provide the perfect building blocks for a novel magnetospinoencephalographic (MSEG) approach to detect neuromagnetic field changes from deep spinal sources and the brain concurrently.

Leveraging these developments in OP-MEG, we have developed a flexible MSEG system that allows for recording spinal and cortical electrophysiology simultaneously. Here we report early and late evoked fields over the cervical spine, in combination with cortical sensory evoked fields (SEFs), in response to median nerve stimulation at the wrist.

## Results

### Concurrent brain, spinal cord, and muscle recordings

Previous limitations in magnetospinography hardware (imposed by cryogenic fixed sensor arrays) have meant that spinal electrophysiology has been recorded in isolation^21,22^. OPM sensors uniquely allow for non-invasive imaging of the brain and spinal cord simultaneously, facilitating electrophysiology recording over the entire human central nervous system (CNS; Figure 1). Here, we chose median nerve stimulation (MNS) to probe the CNS, due to the time-locked nature of the expected response and the wealth of previous literature providing strong priors about the latency of the evoked field^17,21,40,41^.

**Figure 1.**
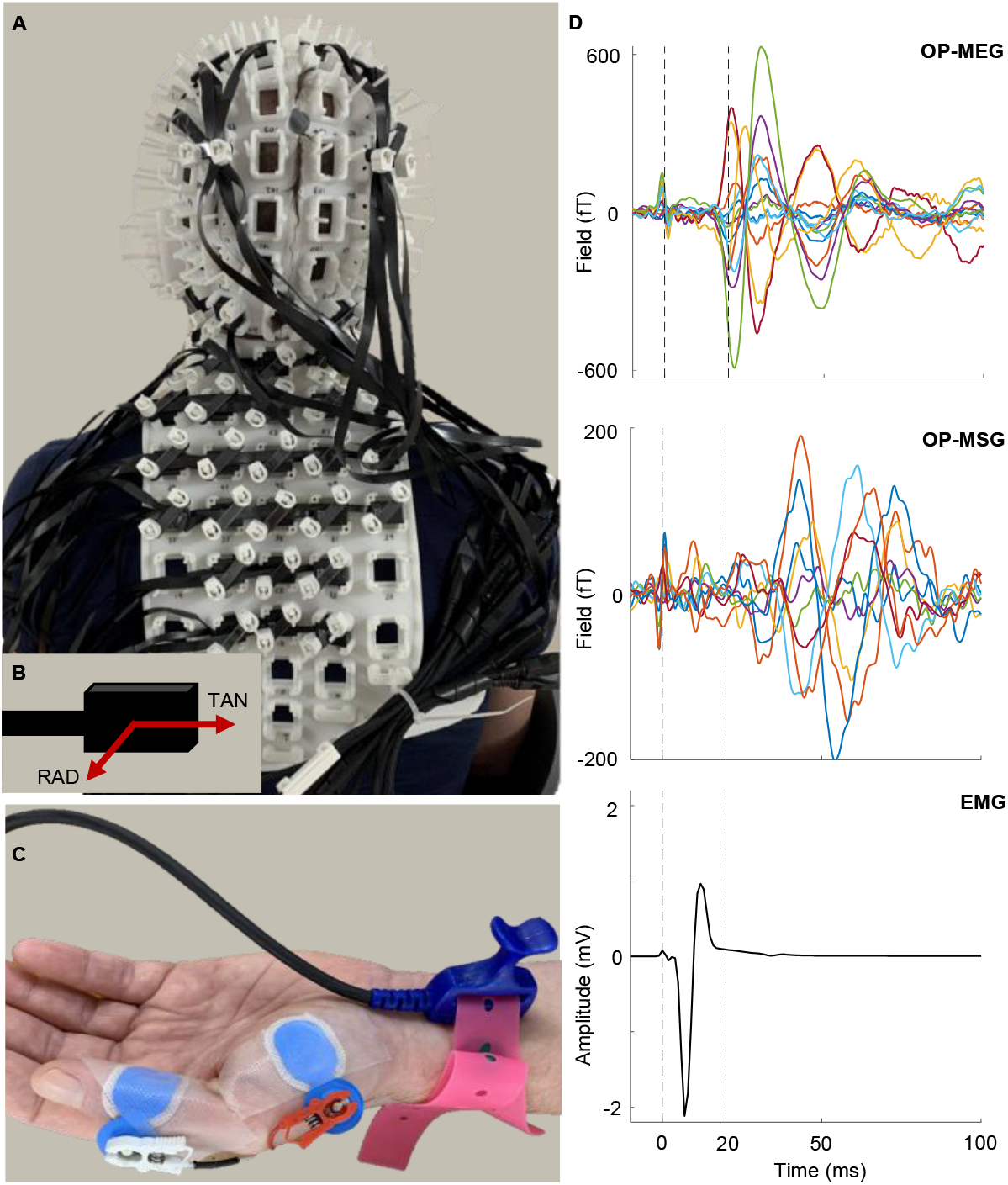
Experimental setup. **A**. 3D printed customised spinal-and head-cast. The head-cast contains optically pumped magnetometer (OPM) sensors over the base of the head and sensorimotor cortex. During the recording the subject is seated on a wooden stool within the magnetically shielded room. **B**. Schematic of OPM sensor, demonstrating the two axes: radial (RAD) and tangential (TAN) to the spinal cord and cortex. **C**. Median nerve stimulation (MNS) electrode applied to the wrist with electromyography (EMG) electrodes recording the abductor pollicis brevis (APB) muscle activity. **D**. Example activity across the nervous system in response MNS, averaged over 2260 trials. From top to bottom: OPM-based magnetoencephalography (OP-MEG) from the left cortical region; OPM-based magnetospinography (OP-MSG); EMG of APB muscle.

We recorded MNS evoked spinal magnetic field changes in 4 participants (A – D). We built one spinal cast optimized to participant A. For this participant we also measured simultaneously from the spinal cord and cortex. In three additional participants (B – D; presented in the supplementary information) we measured evoked spinal magnetic field changes, using the spinal-cast of participant A.

In addition to brain and spinal cord electrophysiology, we recorded electromyography (EMG) from the abductor pollicis brevis muscle, capturing the activity at each level of the hierarchy of the nervous system (Figure 1D). The EMG responses to MNS at the wrist had an onset of 4 ms and peaked 6 – 7 ms post stimulation. Descriptively, we found that the greatest response in global field power recorded at the spinal OPM channels was a 45 – 59 – 72 ms complex, with additional evidence for an early spinal response peaking at 10 ms. Simultaneous cortical evoked fields had an initial global field power peak at 21 ms followed by a 30 ms peak, characteristic of cortical MNS evoked response^42,43^.

### Spinal response to median nerve stimulation

Spinal measurements for participant A took place over four separate sessions, consisting of three right MNS (recorded during sessions 1 – 3) and two left MNS (recorded during session 3 and 4). Magnetic field change in response to MNS was recorded over the surface of the back using a custom-made spinal-cast with 21 – 27 dual-axis sensors, each with two orthogonal measurement channels, oriented anterior-posterior (nominally radial to the cord) and left-right (nominally tangential). Figure 2Ai and ii demonstrate the simulated magnetic field distribution caused by a single spinal source, illustrating the responses one might expect a single current dipole within, and oriented along the axis of, the cervical spinal cord. Field patterns are shown for radially orientated channels (peaking either side of spine with opposite polarity) and tangentially orientated channels (peaking above spine), respectively. For display purposes, in Figure 2Ciii, Biii and Figure 3A we have plotted the spinal cord evoked field (SCEF) time courses for tangentially orientated channels on the midline and the radially orientated channels to the left and right of this (illustrated by Figure 2B), where the largest responses were predicted by the dipole simulation. Conversely, the statistical analysis was conducted on all spinal sensors (both radial and tangential).

**Figure 2.**
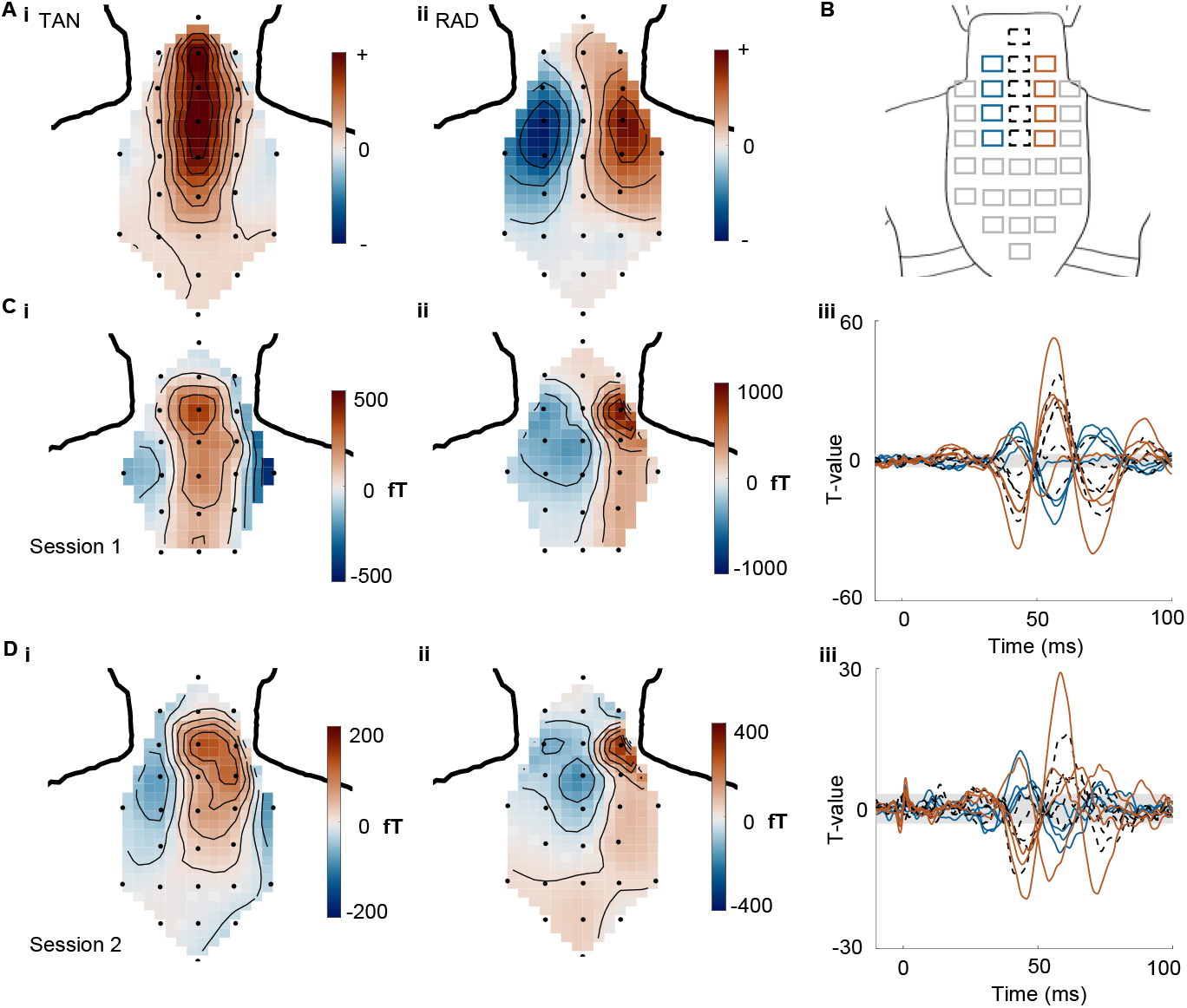
Spinal magnetic field changes in response to right median nerve stimulation (MNS). **A**. Simulated field map of tangentially (i) and radially (ii) orientated channels for a current dipole oriented along the spinal cord. **B**. Sensor locations with colour map identifying the channels plotted in Ciii and Diii. The top four radially orientated channels (solid line) to the left (blue) and right (red) of the midline and five tangentially orientated midline channels (dotted black) are plotted for −10 to 100 ms around stimulus onset. Tangential midline channels were plotted to highlight the central peak activity predicted by the simulated dipole (Ai). Radial channels on the left and right of the midline were plotted in different colours to highlight the change in polarity occurring in the response across the midline predicted by the simulated dipole (Aii). **C**. Magnetic field map of tangentially (i) and radially (ii) orientated channels for 55 – 65 ms post-stimulation and T-values for the spinal cord evoked field (iii) after right wrist MNS during session 1 (averaged over 2260 trials). The grey area represents where the signal is below the false discovery critical height threshold. **D**. As in C but for right wrist MNS during session 2 (averaged over 2260 trials).

**Figure 3.**
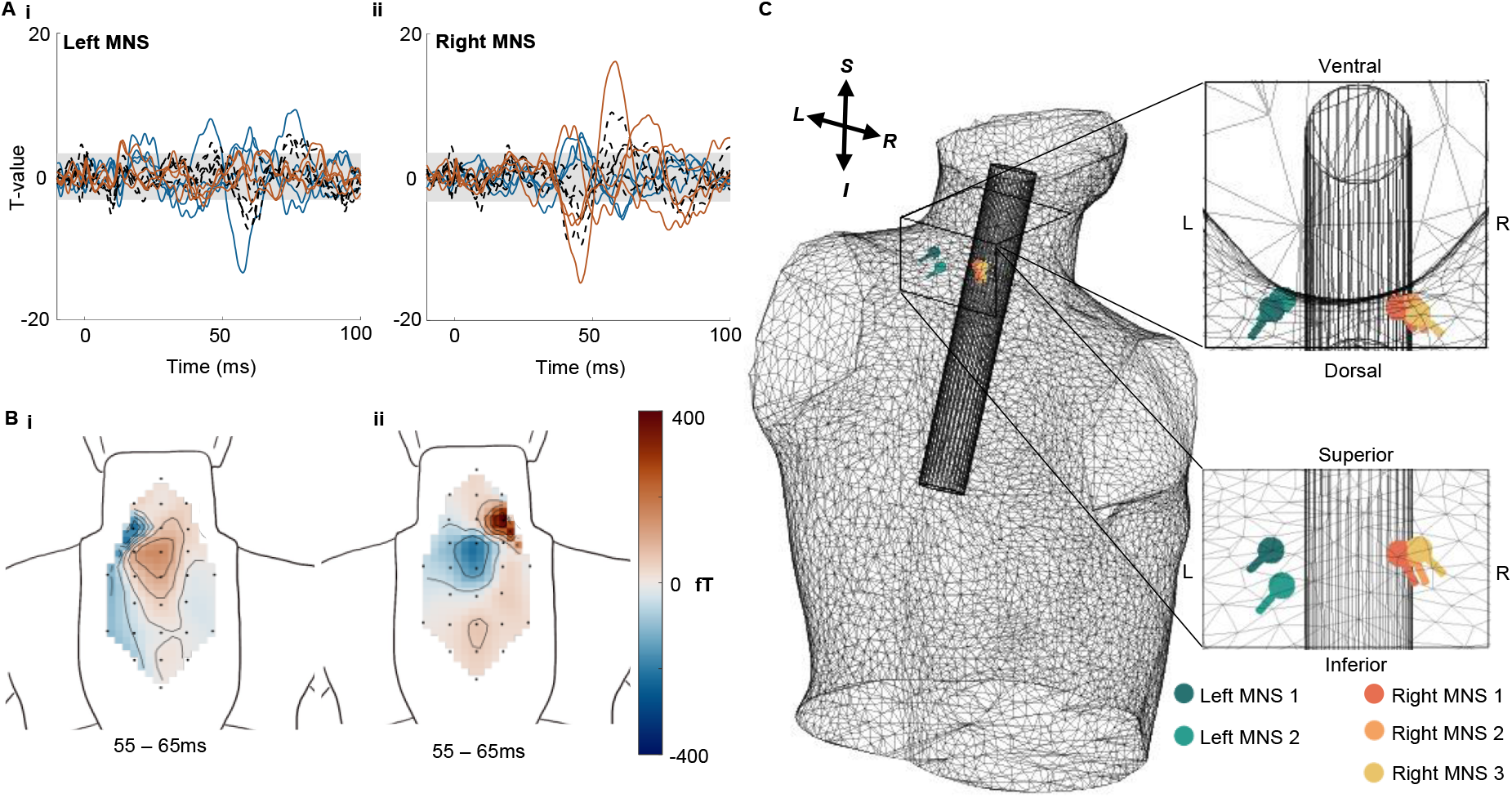
Sensor and source level response to left and right median nerve stimulation (MNS). **A**. T-values for time courses of evoked fields for left (i) and right (ii) MNS during session 3 (each averaged over 1130 trials). Colours indicate channels located to the left (blue, radially orientated), to the right (red, radially orientated), or over the midline (black dotted, tangentially orientated), respectively. The grey area represents where the signal is below the false discovery critical height threshold. **B**. Magnetic field maps for radially oriented channels averaged over 55 – 65 ms post stimulation for left (i) and right (ii) MNS at the wrist. **C**. Optimal dipole fit for the 55 – 65ms component, identified using equivalent current dipole (ECD) fitting for three right MNS sessions (Right MNS 1, 2 and 3; 2260, 2260 and 1130 trials, respectively) and two left MNS sessions (Left MNS 1 and 2; 1130 and 1695 trials, respectively), displayed within the body model relative to a cylinder used to approximate spinal cord location.

During session 1, 2 and 3, 2260, 2260 and 1130 trials, of suprathreshold right wrist MNS were averaged forming the SCEF illustrated in Figure 2Ciii, Diii and Figure 3Aii, respectively. In session 3 and 4, 1130 and 1695 trials of left wrist MNS were averaged to form the SCEF in Figure 3Ai and Supplementary Figure 1Bi. The latency of the peak components of the SCEF were extracted using the peak activity of the absolute maximum T-values across all spinal channels (both radial and tangential) that surpassed the false discovery rate (FDR) threshold. Previous MSG studies have found evoked spinal responses to MNS at the wrist occurring 9.7 ms after stimulation^21^. We found peak activity in the early period (< 12 ms post-stimulation) at 10, 9 and 10 ms, for right MNS in sessions 2 and 3 (Figure 2Dii and 3Aii) and left MNS in session 3 (Figure 3Ai), respectively. Following this, responses at 18, 14, 19, 20 and 19 ms were seen for right wrist MNS session 1 – 3 and left wrist MNS in sessions 3 and 4, respectively (see Supplementary Figure 2 for clearer visualization of early responses).

The highest amplitude spinal OPM response peaked between 55 – 60 ms. Surrounding this central peak are two components with opposite polarity creating a triphasic waveform. The latency of the three peaks of the triphasic waveform, determined by the absolute maximum T-values were 43, 45, 45 ms for peak 1; 56, 58, 58 ms for peak 2; 71, 72, 70 ms for peak 3 for right wrist MNS sessions 1 – 3, respectively, and 45, 38 ms for peak 1; 57, 58 ms for peak 2; 76, 66 ms for peak 3 for left wrist MNS in sessions 3 and 4, respectively. The field maps for the averaged 55 – 65 ms response demonstrate a similar spatial pattern as the simulated dipole, illustrated in Figure 2C and D, for tangential and radial channels following right MNS in session 1 and 2, respectively, and Figure 3Bi and ii for radial channels following left and right MNS in session 3, respectively. The magnetic field changes for the early response, not shown here, were less consistent (field maps in supplementary information).

### Lateralized spinal evoked response for left and right MNS

To determine if the spinal response was dependent on stimulus laterality, during session 3 we recorded MNS at both the left and then right wrist, with 1130 trials each. The averaged magnetic field response to these two conditions can be seen in Figure 3A. We found that left and right MNS exhibited lateralized magnetic field responses (Figure 3A and B), with the largest response (expressed 55 – 60 ms) recorded on the side of the stimulation. This is emphasized by the field maps (Figure 3B) for the radially orientated channels, which highlight how the dipolar pattern differs for left vs right MNS.

### Source analysis of left and right MNS

To further investigate the influence of stimulation side on the spinal evoked field and to interrogate the response in further detail, we conducted a source analysis. We chose to explain the data with an equivalent current dipole (ECD), the simplest possible model. We fit this model to averaged evoked response data (shown in Figure 2C, 2D, 3A and Supplementary Figure 1B) from the three right wrist MNS sessions (labelled Right MNS 1, 2 and 3) and the two left wrist MNS sessions (labelled Left MNS 1 and 2) individually. For the forward model, we used a Nolte single shell model^44^ fit to the participant’s digitized torso (minus head, arms and legs). The non-linear dipole fits were initialized within (but not constrained to) a cylinder used to approximate the spine location. There is clearly a great deal of work to be done on optimal forward and generative models of spinal cord function, however, it was encouraging to see that the optimal dipole fitting for 55 – 65 ms was consistent across sessions and lateralized to side of stimulation (Figure 3C).

## Discussion

We describe a wearable OPM system for the concurrent imaging of electrophysiological signals from the spinal cord and the brain. This study is based on peripheral nerve stimulation but opens up the possibility for non-invasive precise imaging across the entire CNS in humans during natural movement.

Recent work using MSG has described very fast spinal responses occurring 5.5 ms following MNS at the elbow^22^ and 9.7 ms following MNS at the wrist^21^. Here, we saw evidence for early components 10 ms after wrist MNS. These are consistent with the N13 (median nerve) spinal sensory evoked potential that can be recorded with surface electrodes^18^. At this stage, the earliest detectable field changes reported in earlier MSG studies likely stem from the conductivity changes as the depolarization propagates along the spinal nerve, through the intervertebral foramen, to the spinal cord^21^. Subsequent current may reflect the summed synaptic activity and currents from the dorsal column, followed by propagation of the signal along the spinal canal^20,21,40^. Our ability to detect early spinal components may have been restricted by an interaction between the limited bandwidth of our OPM system^45^ and the electrical stimulation artefact. The bandwidth of our system (up to 3dB point) is 135 Hz and is due to the inherent trade-off between bandwidth and sensitivity in zero field OPMs^30,46,47^. As such, the pulse artefact (due to the electrical stimulation) bleeds into early portions of the measured signal. This does not preclude measurement of very fast changes (as the magnitude attenuation due to an effective first order filter is offset by the sensitivity gains from optimal sensor positioning), but it does mean that we need to investigate alternative stimulation or experimental designs (e.g., via collusion experiments) to examine these early components.

In addition to these early responses, we also observed strong subsequent field changes attributed to the spinal cord. These later signal deflections are compatible with early work using an individual SQUID sensor positioned over the spinal cord^41^ and likely reflect a combination of intra-spinal mechanisms, descending feedback signals, and long-latency reflex activity, all from a mixture of anti-and orthodromic stimulation effects of the suprathreshold MNS used here. Long-latency reflexes (LLR) onset at ∼50 ms in thenar EMG and can be elicited by both muscle and cutaneous afferent stimulation^48^. The first peak of the later triphasic spinal component we see here, occurs at ∼40 ms after stimulation, which therefore, may contribute to the LLR. Future work will address the specific contributions of these mechanisms to the signals that can be quantified with MSEG, for which the additional ability to image the brain concurrently will be beneficial.

The electrophysiological responses we report here, at both spinal and cortical levels, have previously been reported and are well-established as markers for clinical assessment^15,16,49,50^. We primarily used peripheral nerve stimulation to establish a proof of principle for the ability of OPMs to detect spinal activity in humans, and to demonstrate the ability for concurrent electrophysiological assessment across the entire CNS. While here initially conducted in a small number of subjects only, this now opens the exciting opportunity to study the spinal responses in a wide range of paradigms in larger cohorts and patient groups.

There are several limitations and challenges we want to point out. The overall coverage of both the back and the head was still relatively limited, owing to the number of OPM sensors available at the time of recording. We have also used simple volume conductor and source models which clearly warrant elaboration. Accurate source reconstruction will be necessary for obtaining images of spinal activity. The specific structure and conductivity profiles of the spinal cord and back create several challenges for this. Going forward, solutions based on boundary/finite element models that will best account for the spinal anatomy are likely to boost precise source estimation, however, we here show that lateralized segmental inference can be obtained even without such approaches.

We expect that with a larger number of sensors, especially by using triaxial rather than dual-axis devices^51,52^, we will be able to better characterize both the signal and environmental noise space. Also, although OPM sensitivity does begin to decline at 130Hz, it does so only as a first order filter (i.e., a halving of amplitude for every doubling of frequency). One limitation of this current study was the stimulus artefact whose impulse response obscured the earlier signal components. Future work could also be devoted to new beamformer schemes (e.g., the recursive null-steering beamformer^53^ implemented in Sumiya et al.^22^ and Akaza et al.^21^) to attenuate these artefacts or indeed simply different experimental paradigms. Despite the current limitations, however, we were able to show lateralized responses to peripheral nerve stimulation, with dipole locations that were compatible with the known anatomy of the spinal cord, and with latencies matching previous reports using surface electrodes or MSG^21,41^.

Additionally, optimizing the orientation and location of the OPM sensors will help to distinguish varying aspects of the spinal response, such as activity in the dorsal root ganglia, intraspinal processing and the ascending wave. Increasing the number of sensors used to provide wider coverage, along with the ongoing reduction in OPM size, will enable high density recordings at a reduced weight, improving the feasibility and tolerability for unconstrained assessment of brain and spinal physiology, and finally, although unlikely to have been an issue here, due to the focus on evoked responses, future recordings may also aim to measure breathing and heart rates to aid in noise reduction techniques.

In the present study we chose to use peripheral stimulation to probe the sensorimotor system, which benefits from the time-locked nature of the response. Future progressions of this research should study the cortico-spinal interactions of movement generation, but this is not without challenges. Due to the multiple stages in motor processing before the spinal cord, activity may be more temporally dispersed, potentially leading to weaker signals. Additionally, movement artefacts will create challenges in data processing, however, previous OP-MEG work has identified analysis pipelines to overcome this^34,54^. Moreover, changes in power may provide greater insight into motor processing at the spinal level, which along with OP-MEG recordings could illuminate cortico-spinal interactions^55^.

One major advance of MSEG over existing approaches to spinal electrophysiological recordings is that these sensors can be worn, offering widespread usability to study the electrophysiology of the spinal cord not only in healthy adults but a range of cohorts such as children or many disorders characterized by damaged or pathological movement^32,56^. The potential for studying movement is particularly beneficial in patient groups where remaining still is not possible or movement is impaired. One of the most transformative aspects of OPM recordings is that high quality recordings can be carried out in a variety of positions of the subject, even whilst subjects make relatively large and unpredictable movements^29,32–34,57^, meaning that OP-MEG can be used in a wide range of postures and to study participants groups that may not remain still. Therefore, this technology lends itself to the study of motor development as well as pathology in conditions such as stroke, Parkinson’s disease and neurogenerative disorders. The flexibility of OPM sensor arrangement also means that sensor arrays are not limited to the cervical spinal cord, as done here, but can easily be configured for thoracic and lumbosacral spinal imaging, or indeed to image the entire spinal cord, whilst additionally allowing for recording brain activity with high precision.

A wide range of disorders and pathologies affect the spinal cord and cortico-spinal interplay, including spinal cord injury (SCI), stroke, traumatic brain injury, or the spinal grey matter degeneration seen in amyotrophic lateral sclerosis^8,58–60^. Basic understanding of the underlying mechanisms of the impairments caused by these pathologies, the response to treatment, and basis of recovery will benefit from imaging the entire CNS. The ability for high-precision electrophysiology of the entire CNS will transform our understanding of the human spinal cord.

## Methods

This study was approved by the University College London research ethics committee. Four participants (two male) took part in the present study after providing informed written consent. The initial recordings were conducted on one male, labelled participant A (aged 45) for whom the scanner cast was customised for (see below for more detail). Participant A underwent four separate testing sessions. Sessions 2 and 3 took place on consecutive days, 1 month after session 1. Session 4 occurred 7 months after session 1. An additional three participants, labelled B, C and D (aged 26, 28 and 31, respectively) completed one session each using the scanner cast customised for participant A. These data are reported in the supplementary information.

### Experimental setup

#### Scanner cast design and development

An optical white light scanner was used to generate a 3D model of participant A’s upper body for which the spinal OPM scanner cast was created (Chalk Studios, London, UK). The spinal scanner cast, providing coverage of the neck and back, was 3D printed in nylon with Velcro straps attached to hold the cast in place during scanning (Figure 1A). The spinal scanner cast contains a total of 33 OPM sensor slots, of which a maximum of 27 slots were utilized in the current study. For the cortical sensor array a generic head-cast was used to capture the magnetic field over the contralateral sensorimotor cortex.

#### Data acquisition (OPM recording)

All data were recorded in a magnetically shielded room (MSR), with internal dimensions of 438 × 338 × 218 cm, which included several design optimizations to improve magnetic shielding (Magnetic Shields Ltd, Staplehurst, UK). The room comprises of two inner layers of 1mm mu-metal, a 6mm copper layer and two external layers of 1.5mm mu-metal. Magnetic equilibration of the MSR was accomplished prior to each scanning session through degaussing, during which a sinusoidal current with decreasing amplitude is applied to coils wound around the shielding material inducing a magnetic flux in a closed loop^61^. Automated sensor-level nulling with inbuilt field cancellation coils prior to every experimental recording further reduces residual fields around the vapour cell of the OPM sensors^34,62^. Two reference OPM sensors were placed away from the participant, in fixed positions within the MSR, to capture the environmental magnetic field changes. During the experiment, data were recorded from the OPM sensors at a sampling rate of 6000 Hz from both axes, which were radial and tangential to the measurement surface (either back or head). Here we use the term ‘sensor’ to refer to the physical device and the term ‘channel’ to refer to either the radially or tangentially oriented axis within the sensor. Data were saved in brain imaging data structure (BIDS) format.

#### Recording sessions

Participant A underwent four separate recording sessions. An array consisting of 21, 27, 27 and 24 dual-axis OPM sensors (QuSpin Inc., Louisville, USA) in session 1 – 4 respectively were arranged in the custom-made spinal scanner cast to cover the neck and upper back. Variability in sensor number was due to sensor availability at the time of recording and differences in sensor arrangement occurred at the edges of the array to ensure the cervical spinal cord was always well covered. In addition to the spinal coverage, session 2 and 3, also comprised cortical recordings from 15 OPM sensors that were arranged over the left sensorimotor cortex in a generic 3D printed OPM helmet. The OPM sensors have a sensitivity of ∼15 fT/√Hz between 10 and 100 Hz.

#### Median nerve stimulation (MNS)

The median nerve was stimulated at the wrist via a peripheral nerve stimulation electrode with the cathode positioned distally to the anode (Figure 1C). Pulses were delivered with a 50 μs pulse-width at a stimulation frequency of 2Hz using a DS7A current stimulator (Digitimer). A total of 565 stimulations were delivered for each run with a short break between runs. Stimulation was supramaximal to induce a thumb twitch (stimulator intensity 12 – 31.2 mA). For participant A, session 1 and session 2 consisted of four runs of right MNS (a total of 2260 trials). Session 3 consisted of two runs of each right and left MNS (1130 per wrist) and session 4 consisted of nine runs of left MNS (three peri-motor threshold, three 1.5 x threshold and three 2.5 x threshold; 1695 trials each) and three cutaneous (next to the nerve) stimulation runs (1695 trials), see supplementary information for further details. Four runs were recorded for participant B and C for the right wrist and for participant D for the left wrist (a total of 2260 per participant).

### Dipole simulation

A dipole simulation was conducted in FieldTrip software^63^ to visualize the expected change in magnetic field recorded by the spinal sensor array. The sensor locations from the spinal array in session 2 (total of 27 sensors) were used to conduct the dipole simulation. A single shell model^44^ was created using the 3D optical scan of the participants torso (with the arms and head removed). Within this model, the dipole was oriented upwards towards the head, and placed at the estimated location of the spine (level to the middle of the shoulders). The FieldTrip function `ft_dipolesimulation` was used to project the simulated dipole field back onto the sensor locations. The field map for the simulated sensor data was then plotted on the map of the radially and tangentially orientated channels of the spinal sensors.

### Data analysis

#### Data pre-processing

All data pre-processing was conducted in MATLAB R2020b (Mathworks Inc., Natick, MA, USA) using the statistical parameter mapping (SPM) toolbox. The OPM BIDS formatted data were loaded into SPM format and filtered using homogenous field correction^64^. A 2^nd^ order bidirectional Butterworth bandpass filter between 20 – 300 Hz was used to remove low and high frequency noise. Next, in order to reduce external noise synthetic gradiometry was conducted^65^, where the linear regression of the two reference sensors was subtracted from the spinal and cortical sensor data. Synthetic gradiometry was additionally done with narrow-band reference sensor data, bandpass filtered between both 47 – 53 Hz and 97 – 103 Hz, to reduce line noise. Synthetic gradiometry was chosen over a notch filter for line noise reduction to minimize the stimulation artefact distortion caused by filtering. Individual trials were epoched from −100 to +300 ms around stimulus onset and baseline corrected to the average signal between −100 to 0 ms. Trials from subsequent runs of the same task within sessions were merged and averaged.

#### Statistics

T-statistics were computed for the three right MNS sessions and two left MNS sessions, independently. A student’s T-Test was calculated on the SCEF time course, from −10 to +100 ms around stimulation, for all spinal channels (both radial and tangential). The false discovery critical height threshold was calculated using the spm_uc_FDR function to correct for multiple comparisons. For this, the false discovery rate was set to 0.01 (0.05 divided by five sessions).

#### Source localization

Equivalent current dipole (ECD) fitting was used to find the optimal dipole location and orientation for the 55 – 65 ms response, using FieldTrip software^63^. The window of 55 – 65 ms is in line with the central peak of the spinal 43-58-89 ms complex evoked following wrist MNS, as previously described by Mizutani and Kuriki^41^. Trial averaged left and right MNS SCEFs were averaged in the 55 – 65ms window before dipole fitting was separately completed for each session.

To produce a volume conductor model, the optical 3D scan, used to generate the spinal scanner cast, was converted into a mesh. The arms and head were removed to produce an empty torso single shell model used for the forward model^44^. A cylinder (diameter 40 mm), within the upper back of the torso model, was used to approximate the location of the spinal cord. Once the optimal dipole fit was established within the cylinder grid, a non-linear search was conducted, constrained within the torso model. This identified the optimal coordinate and orientation of the 55 – 65ms dipole.

## Acknowledgments

This work was initiated (and T.M.T. funded) through a Wellcome collaborative award (203257/Z/16/Z, 203257/B/16/Z) and is now funded through a Wellcome Technology development award (223736/Z/21/Z). G.C.O. was funded through EPSRC (EP/T001046/1) funding from the Quantum Technology hub in sensing and timing (sub-award QTPRF02). L.C.M. was funded through Medical Research Council (MR/N013867/1). C.Z. was funded through Brain Research UK (201718-13). The Wellcome Centre for Human Neuroimaging is supported by core funding from the Wellcome Trust (203147/Z/16/Z). We would also like to acknowledge the support of Mark Lim of Chalk Studios in designing the 3D printed scanner-casts used in this study. We would like to thank Dr Adachi and Professor Kawabata from the Kanazawa Institute of Technology and Tokyo Medical and Dental University in Japan for their support and advice in the conception of this project.

## References

1. Filli, L. & Schwab, M. E. The rocky road to translation in spinal cord repair. Annals of Neurology vol. 72 (2012).

2. Giove, F. et al. Issues about the fMRI of the human spinal cord. in Magnetic Resonance Imaging vol. 22 (2004).

3. Dauleac, C., Frindel, C., Mertens, P., Jacquesson, T. & Cotton, F. Overcoming challenges of the human spinal cord tractography for routine clinical use: a review. Neuroradiology vol. 62 (2020).

4. Stroman, P. W. et al. The current state-of-the-art of spinal cord imaging: Methods. NeuroImage vol. 84 (2014).

5. Furby, J. et al. Magnetic resonance imaging measures of brain and spinal cord atrophy correlate with clinical impairment in secondary progressive multiple sclerosis. Multiple Sclerosis 14, (2008).

6. Freund, P. et al. Degeneration of the Injured Cervical Cord Is Associated with Remote Changes in Corticospinal Tract Integrity and Upper Limb Impairment. PLoS ONE 7, (2012).

7. Panara, V. et al. Correlations between cervical spinal cord magnetic resonance diffusion tensor and diffusion kurtosis imaging metrics and motor performance in patients with chronic ischemic brain lesions of the corticospinal tract. Neuroradiology 61, (2019).

8. Grabher, P. et al. Tracking sensory system atrophy and outcome prediction in spinal cord injury. Annals of Neurology 78, (2015).

9. Sprenger, C., Finsterbusch, J. & Büchel, C. Spinal cord-midbrain functional connectivity is related to perceived pain intensity: A combined spino-cortical fMRI study. Journal of Neuroscience 35, (2015).

10. Baliki, M. N., Baria, A. T. & Vania Apkarian, A. The Cortical Rhythms of Chronic Back Pain. Journal of Neuroscience 31, (2011).

11. Moore, K. A. et al. Partial peripheral nerve injury promotes a selective loss of GABAergic inhibition in the superficial dorsal horn of the spinal cord. Journal of Neuroscience 22, (2002).

12. Lamy, J. C. et al. Impaired efficacy of spinal presynaptic mechanisms in spastic stroke patients. Brain 132, (2009).

13. Karbasforoushan, H., Cohen-Adad, J. & Dewald, J. P. A. Brainstem and spinal cord MRI identifies altered sensorimotor pathways post-stroke. Nature Communications 10, (2019).

14. Urasaki, E. et al. Skin and epidural recording of spinal somatosensory evoked potentials following median nerve stimulation: correlation between the absence of spinal N13 and impaired pain sense. Journal of Neurology 237, 410–415 (1990).

15. Prestor, B., Žgur, T. & Dolenc, V. v. Subpially recorded cervical spinal cord evoked potentials in syringomyelia. Electroencephalography and Clinical Neurophysiology/ Evoked Potentials 80, (1991).

16. Imajo, Y. et al. Assessment of spinal cord relative vulnerability in C4–C5 compressive cervical myelopathy using multi-modal spinal cord evoked potentials and neurological findings. The Journal of Spinal Cord Medicine 44, 541–548 (2021).

17. Cracco, R. Q. Spinal evoked response: Peripheral nerve stimulation in man. Electroencephalography and Clinical Neurophysiology 35, (1973).

18. Restuccia, D., Valeriani, M., di Lazzaro, V., Tonali, P. & Mauguiere, F. Somatosensory evoked potentials after multisegmental upper limb stimulation in diagnosis of cervical spondylotic myelopathy. Journal of Neurology Neurosurgery and Psychiatry 57, (1994).

19. Fujimoto, H. et al. Differential recording of upper and lower cervical N13 responses and their contribution to scalp recorded responses in median nerve somatosensory evoked potentials. Journal of the Neurological Sciences 187, (2001).

20. Iragui, V. J. The cervical somatosensory evoked potential in man: Far-field, conducted and segmental components. Electroencephalography and Clinical Neurophysiology 57, (1984).

21. Akaza, M. et al. Noninvasive measurement of sensory action currents in the cervical cord by magnetospinography. Clinical Neurophysiology 132, 382–391 (2021).

22. Sumiya, S. et al. Magnetospinography visualizes electrophysiological activity in the cervical spinal cord. Scientific Reports 7, 2192 (2017).

23. Kawabata, S., Komori, H., Mochida, K., Harunobu, O. & Shinomiya, K. Visualization of conductive spinal cord activity using a biomagnetometer. Spine (Phila Pa 1976) 27, (2002).

24. Adachi, Y. et al. Magnetospinography: Instruments and Application to Functional Imaging of Spinal Cords. IEICE Transactions on Electronics E 96.C, 326–333 (2013).

25. Ushio, S. et al. Visualization of the electrical activity of the cauda equina using a magnetospinography system in healthy subjects. Clinical Neurophysiology 130, (2019).

26. Sakaki, K. et al. Evaluation of neural activity by magnetospinography with 3D sensors. Clinical Neurophysiology 131, (2020).

27. Singh, S. P. Magnetoencephalography: Basic principles. Ann Indian Acad Neurol 17, (2014).

28. Miyano, Y. et al. Visualization of electrical activity in the cervical spinal cord and nerve roots after ulnar nerve stimulation using magnetospinography. Clinical Neurophysiology 131, 2460–2468 (2020).

29. Boto, E. et al. Moving magnetoencephalography towards real-world applications with a wearable system. Nature 555, 657–661 (2018).

30. Tierney, T. M. et al. Optically pumped magnetometers: From quantum origins to multi-channel magnetoencephalography. NeuroImage vol. 199 (2019).

31. Boto, E. et al. A new generation of magnetoencephalography: Room temperature measurements using optically-pumped magnetometers. Neuroimage 149, 404–414 (2017).

32. Hill, R. M. et al. A tool for functional brain imaging with lifespan compliance. Nature Communications 10, (2019).

33. Roberts, G. et al. Towards OPM-MEG in a virtual reality environment. Neuroimage 199, (2019).

34. Seymour, R. A. et al. Using OPMs to measure neural activity in standing, mobile participants. Neuroimage 244, (2021).

35. Iivanainen, J., Stenroos, M. & Parkkonen, L. Measuring MEG closer to the brain: Performance of on-scalp sensor arrays. Neuroimage 147, (2017).

36. Boto, E. et al. On the potential of a new generation of magnetometers for MEG: A beamformer simulation study. PLoS ONE 11, (2016).

37. Lin, C. H. et al. Using optically pumped magnetometers to measure magnetoencephalographic signals in the human cerebellum. Journal of Physiology 597, (2019).

38. Tierney, T. M. et al. Mouth magnetoencephalography: A unique perspective on the human hippocampus. Neuroimage 225, 117443 (2021).

39. Barry, D. N. et al. Imaging the human hippocampus with optically-pumped magnetoencephalography. Neuroimage 203, (2019).

40. Jones, S. J. Short latency potentials recorded from the neck and scalp following median nerve stimulation in man. Electroencephalography and Clinical Neurophysiology 43, 853–863 (1977).

41. Mizutani, Y. & Kuriki, S. Somatically Evoked Magnetic Fields in the Vicinity of the Neck. IEEE Transactions on Biomedical Engineering BME-33, (1986).

42. Allison, T., Mccarthy, G., Wood, C. C. & Jones, S. J. Potentials evoked in human and monkey cerebral cortex by stimulation of the median nerve: A review of scalp and intracranial recordings. Brain 114, (1991).

43. Kakigi, R. Somatosensory evoked magnetic fields following median nerve stimulation. Neuroscience Research 20, (1994).

44. Nolte, G. The magnetic lead field theorem in the quasi-static approximation and its use for magnetoenchephalography forward calculation in realistic volume conductors. Physics in Medicine and Biology 48, (2003).

45. Marquetand, J. et al. Optically pumped magnetometers reveal fasciculations non-invasively. Clinical Neurophysiology 132, (2021).

46. Cohen-Tannoudji, C., Dupont-Roc, J., Haroche, S. & Laloë, F. Diverses résonances de croisement de niveaux sur des atomes pompés optiquement en champ nul. I. Théorie. Revue de Physique Appliquée 5, (1970).

47. Dupont-Roc, J., Haroche, S. & Cohen-Tannoudji, C. Detection of very weak magnetic fields (10-9gauss) by 87Rb zero-field level crossing resonances. Physics Letters A 28, (1969).

48. Deuschl, G. & Lücking, C. H. Physiology and clinical applications of hand muscle reflexes. Electroencephalography and clinical neurophysiology. Supplement vol. 41 (1990).

49. Curt, A. & Dietz, V. Traumatic cervical spinal cord injury: Relation between somatosensory evoked potentials, neurological deficit, and hand function. Archives of Physical Medicine and Rehabilitation 77, (1996).

50. Cruse, D., Norton, L., Gofton, T., Young, G. B. & Owen, A. M. Positive Prognostication from Median-Nerve Somatosensory Evoked Cortical Potentials. Neurocritical Care 21, (2014).

51. Brookes, M. J. et al. Theoretical advantages of a triaxial optically pumped magnetometer magnetoencephalography system. Neuroimage 236, (2021).

52. Tierney, T. M. et al. Spherical harmonic based noise rejection and neuronal sampling with multi-axis OPMs. bioRxiv 2021.12.22.473837 (2021).

53. Kumihashi, I. & Sekihara, K. Array-gain constraint minimum-norm spatial filter with recursively updated gram matrix for biomagnetic source imaging. IEEE Transactions on Biomedical Engineering 57, (2010).

54. Seymour, R. A. et al. Interference suppression techniques for OPM-based MEG: Opportunities and challenges. Neuroimage 247, 118834 (2022).

55. Oya, T., Takei, T. & Seki, K. Distinct sensorimotor feedback loops for dynamic and static control of primate precision grip. Communications Biology 3, (2020).

56. Vivekananda, U. et al. Optically pumped magnetoencephalography in epilepsy. Annals of Clinical and Translational Neurology 7, (2020).

57. Holmes, N. et al. A bi-planar coil system for nulling background magnetic fields in scalp mounted magnetoencephalography. Neuroimage 181, (2018).

58. Chen, X. et al. Abnormal functional corticomuscular coupling after stroke. NeuroImage: Clinical 19, (2018).

59. Nagamoto-Combs, K., Morecraft, R. J., Darling, W. G. & Combs, C. K. Long-term gliosis and molecular changes in the cervical spinal cord of the rhesus monkey after traumatic brain injury. Journal of Neurotrauma 27, (2010).

60. Cohen-Adad, J. et al. Involvement of spinal sensory pathway in ALS and specificity of cord atrophy to lower motor neuron degeneration. Amyotrophic Lateral Sclerosis and Frontotemporal Degeneration 14, (2013).

61. Altarev, I. et al. Minimizing magnetic fields for precision experiments. Journal of Applied Physics 117, (2015).

62. Osborne, J., Orton, J., Alem, O. & Shah, V. Fully integrated, standalone zero field optically pumped magnetometer for biomagnetism. in (2018).

63. Oostenveld, R., Fries, P., Maris, E. & Schoffelen, J. M. FieldTrip: Open source software for advanced analysis of MEG, EEG, and invasive electrophysiological data. Computational Intelligence and Neuroscience 2011, (2011).

64. Tierney, T. M. et al. Modelling optically pumped magnetometer interference in MEG as a spatially homogeneous magnetic field. Neuroimage 244, (2021).

65. Fife, A. A. et al. Synthetic gradiometer systems for MEG. IEEE Transactions on Applied Superconductivity 9, (1999).

