## Supplementary information for "Concurrent spinal and brain imaging with optically pumped magnetometers"

### Methods

#### *Varying median nerve stimulation intensity*

During session 4, the stimulation intensity was set to peri-motor threshold (MT), 1.5 x MT and 2.5 x MT. Three runs of 565 trials were recorded at each intensity, resulting in a total of 1695 runs each. In addition to this, to determine whether magnetic field changes at the spinal cord could be induced by sensory stimulation of the skin at the wrist, the stimulating electrodes were moved laterally off the median nerve so that they were just stimulating the skin with the same stimulation intensity as 2.5 x MT. Three runs were recorded for this control condition, resulting in a total of 1695 trials.

#### *Additional participants*

The additional three participants (B – D) that took part in this study used the same spinal scanner cast as participant A, without the head-cast. Due to the personalisation of the scanner cast, the cast was not an optimal fit for these additional participants but provided some cervical spine coverage. For participant B, C and D purely cervical spinal recordings were carried out using 14, 14 and 21 OPM sensors respectively. Spinal cord evoked fields (SCEFs) were recorded for right wrist MNS, over four runs (2260 trials), for participant B and C, and left wrist MNS, over four runs (2260 trials), for participant D.

### Results

#### *Varying spinal cord evoked fields with increasing intensity*

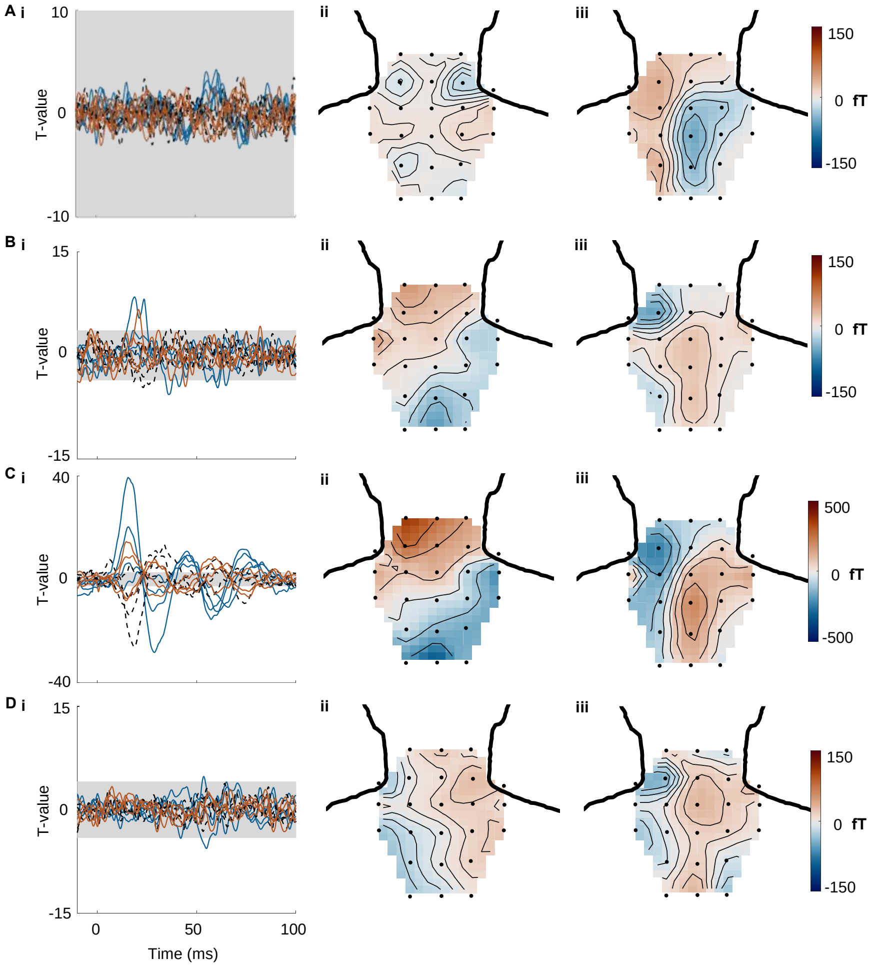


**Supplementary Figure 1.** Spinal cord magnetic field changes to left wrist MNS over different stimulation conditions for participant A. **A.** peri-, **B.** 1.5 x and **C.** 2.5 x motor threshold (MT), and **D.** 2.5 x MT stimulator intensity when applied to the left wrist but not over median nerve. For each condition spinal cord evoked fields (SCEFs) from -10 to +100 around stimulation for radially orientated left and right channels and tangentially orientated middle channels. The grey area represents where the signal is below the false discovery critical height threshold. In Ai, the threshold for which the critical expected false discovery rate (0.01) is controlled for could not be determined (i). Magnetic field maps for radially orientated channels averaged over 10 – 20ms (ii) and magnetic field maps for radially orientated channels averaged over 55 – 65 ms post stimulation (iii) are illustrated. Average of 1695 trials for each condition.

#### *Early spinal cord evoked response*

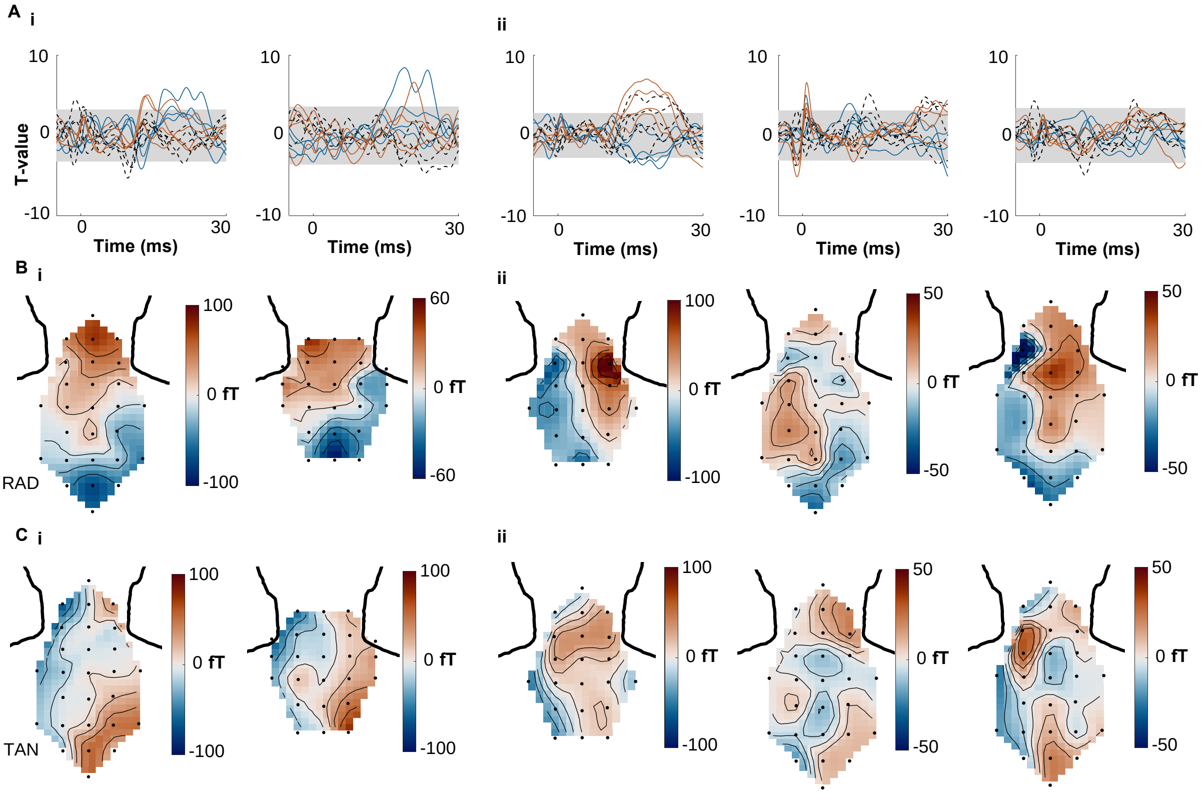


**Supplementary Figure 2.** Early spinal cord magnetic field changes to left and right median nerve stimulation (MNS) for participant A. **A.** T-values for spinal cord evoked fields (SCEFs) to two left (i) and three right wrist (ii) MNS conditions, averaged over, 1130, 1695, 2260, 2260 and 1130 trials, respectively. Radially orientated channels (solid line) to the left (blue) and right (red) of the midline and tangentially orientated midline channels (dotted black) are plotted for -5 to 30 ms around stimulus onset. The vertical dotted lines mark 0 and 20 ms around stimulation. The grey area represents where the signal is below the false discovery critical height threshold. **B.** Magnetic field map of radially orientated channels from 10 to 20 ms post stimulation for left (i) and right (ii) MNS sessions. Different sensor numbers were due to the availability of OPM sensors at the time of recording. **C.** Similar to B but for tangentially orientated channels.

***Additional participants***

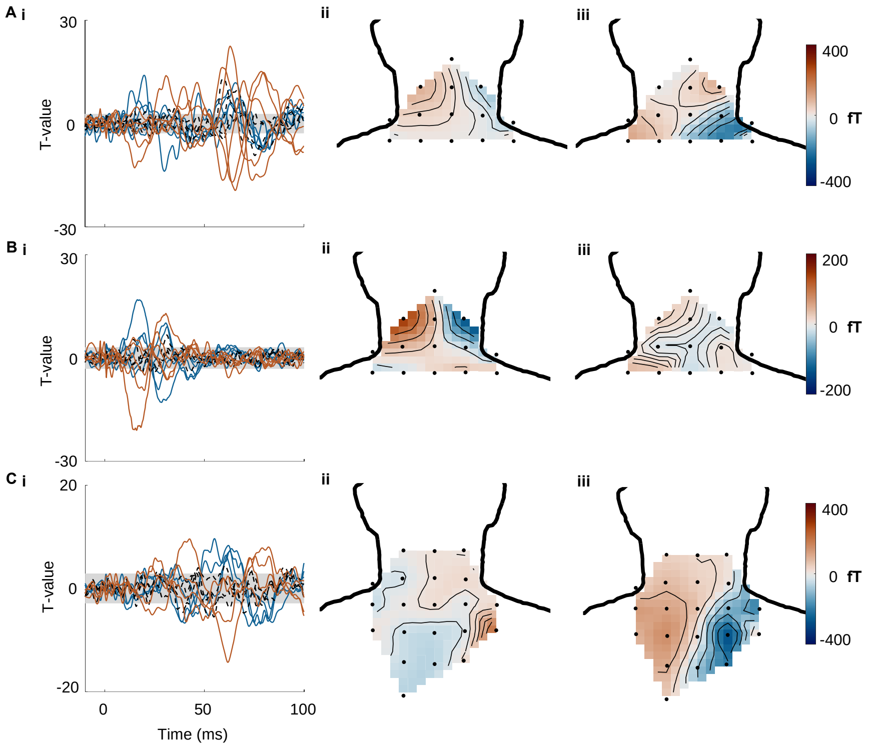


**Supplementary Figure 3.** Magnetic field changes to median nerve stimulation (MNS) for participants B – D. **A.** T-values for spinal cord evoked field (SCEF) to right wrist MNS for participant B (i). Radially orientated channels (solid line) to the left (blue) and right (red) of the midline and tangentially orientated midline channels (dotted black) are plotted for -10 to 100 ms around stimulus onset. The grey area represents where the signal is below the false discovery critical height threshold. Magnetic field map for 10 – 20 ms (ii) and 55 – 65 ms (iii) post stimulation. **B.** As in A for participant C. **C.** Similar to A and B but for left wrist MNS in participant D. Data averaged over 2260 trials for each participant. Different sensor numbers were due to the availability of OPM sensors at the time of recording.
